# Increased expression of viral sensor MDA5 in pancreatic islets and in hormone-negative endocrine cells in recent onset type 1 diabetic donors

**DOI:** 10.1101/2021.11.29.470364

**Authors:** Laura Nigi, Noemi Brusco, Giuseppina E. Grieco, Daniela Fignani, Giada Licata, Caterina Formichi, Elena Aiello, Lorella Marselli, Piero Marchetti, Lars Krogvold, Knut Dahl Jorgensen, Guido Sebastiani, Francesco Dotta

## Abstract

The interaction between genetic and environmental factors determines the development of type 1 diabetes (T1D). Some viruses are capable of infecting and damaging pancreatic β-cells, whose antiviral response could be modulated by specific viral RNA receptors and sensors such as Melanoma Differentiation Associated gene 5 (MDA5), encoded by the IFIH1 gene. MDA5 has been shown to be involved in pro-inflammatory and immunoregulatory outcomes, thus determining the response of pancreatic islets to viral infections. Although the function of MDA5 has been previously well explored, a detailed immunohistochemical characterization of MDA5 in pancreatic tissues of non-diabetic and T1D donors is still missing. In the present study we used multiplex immunofluorescence imaging analysis to characterize MDA5 expression and distribution in pancreatic tissues obtained from 22 organ donors (10 non-diabetic autoantibody-negative, 2 autoantibody-positive, 8 recent-onset and 2 long-standing T1D).

In non-diabetic control donors, MDA5 was expressed both in α- and in β-cells. The colocalization rate imaging analysis showed that MDA5 was preferentially expressed in α-cells.

In T1D donors, we observed an increased colocalization rate MDA5-glucagon respect to MDA5-insulin in comparison to non-diabetic controls; such increase was more pronounced in recent onset respect to long standing T1D donors. Of note, an increased colocalization rate MDA5-glucagon was found in insulin-deficient-islets (IDI) respect to insulin containing islets (ICI).

Strikingly, in T1D donors we detected the presence of MDA5-positive/hormones-negative endocrine islet-like clusters, putatively deriving from dedifferentiation or neogenesis phoenomena. These clusters were exclusively identified in recent onset donors and not detected in autoantibody-positive non-diabetic or T1D long-standing ones.

In conclusion, we showed that MDA5 is preferentially expressed in α-cells and its expression is increased in recent onset T1D donors. Finally, we observed that MDA5 may also characterize the phenotype of dedifferentiated or newly forming islet cells, thus opening to novel roles for MDA5 in pancreatic endocrine cells.

## 1 Introduction

Type 1 diabetes mellitus (T1D) is a chronic autoimmune disease characterized by the progressive destruction of pancreatic β-cells by the immune system, leading to the absolute loss of insulin secretory function. Although molecular mechanisms involved in T1D pathogenesis are still under investigation, disease progression and onset have been widely recognized as the result of the interaction between a predisposing genetic background and environmental factors which may trigger β-cells destruction and immune system activation (1). Among identified environmental factors, viral infections have been undoubtedly shown to play a prominent role (2,3). As a matter of fact, multiple evidence of *Enterovirus*infection of pancreatic islets in T1D donors have now been confirmed by several studies, also adopting rigorous methodological cross-validation approaches in various patients cohorts (4–10).

β-cells susceptibility to viral infections and their innate immune signaling activation upon virus entry are key determinants of subsequent islet inflammatory response (11). Indeed, we and others have previously shown that one of the isoforms of the main Coxsackieviruses entry receptor, namely Coxsackie Adenovirus Receptor (CAR), is preferentially expressed in β-cells in human pancreas, thus conferring specificity and vulnerability of β-cells to certain viruses (12). In addition, it has been recently confirmed that the expression of several markers of interferon (IFN) signature (e.g. MxA, PKR, and HLA-I) in pancreatic islets of T1D and of islet autoantibody-positive donors, is tightly correlated with the presence of enteroviral capsid protein-1 (VP1), thus showing the existence of an antiviral machinery actively contributing to the islet inflammatory response during viral infections in T1D (13). Of note, antiviral signaling molecules initiating downstream inflammatory pathways activation, have been previously detected in human pancreatic islets and linked to the pathogenesis of fulminant T1D (14). Indeed, the activation of antiviral signaling mechanisms are initiated by specific intracellular sensors of viral nucleic acids and/or components. Among such sensors, the melanoma differentiation-associated gene 5 (MDA5) is of utmost importance. MDA5, part of the retinoic acid inducible gene 1 (RIG-I)-like receptor (RLR) family and encoded by the IFIH1 gene, is a dsRNA binding protein which preferentially recognizes viral intermediates long dsRNAs through the interaction with its helicase domain (15). As such, MDA5 is required for intracellular immunity against several classes of viruses, including Picornaviruses family and, consequently, the Enteroviruses Genus. Once activated, MDA5 leads to the downstream recruitment of mitochondrial antiviral signaling protein (MAVS) and the subsequent triggering of a signaling cascade culminating in the activation of nuclear factor‐kappa B (NFκB), interferon‐regulating factor IRF3 and IRF7 (15). Notably, MDA5 knockdown in β-cells decreased dsRNA-induced cytokines and chemokines expression, thus limiting the inflammatory response (16).

The importance of MDA5 function in T1D pathogenesis is also highlighted by the existence of several SNPs conferring increased risk or protection from T1D (17–19). Polymorphisms conferring increased risk of T1D (e.g. A946T, TT risk genotype) have been shown to induce a weaker interferon-mediated inflammatory-response in human pancreatic islets infected with Coxsackieviruses, confirming the role of MDA5 in antiviral inflammatory response and the effects of these polymorphisms on its function (20). As such, a reduced function of MDA5 may lead to a defective viral clearance and long-lasting virus persistence, thus causing a mild inflammation and potential activation of autoimmunity in genetically susceptible individuals.

Although previous studies started to investigate the role of MDA5 in β-cells (20,21) and of its activation upon viral infection or following inflammatory stresses, both in rodents and in man (14,20–23), a full characterization of its expression and distribution in human pancreatic tissues both in T1D and in non-pathological conditions has not yet been analyzed in detail.

Against this background, we presently analyzed MDA5 expression distribution using multiplex immunofluorescence technique in human pancreatic tissues obtained from non-diabetic, autoantibody-positive and T1D donors from three different organ donor biorepositories (EUnPOD, nPOD and DiViD study), uncovering qualitative and quantitative differences in the pattern of MDA5 expression occurring in T1D vs non diabetic pancreas.

## 2 Materials and Methods

### 2.1 Donors

Formalin-fixed and paraffin embedded (FFPE) human pancreatic sections were obtained from n=10 non-diabetic autoantibody-negative, n=2 autoantibody-positive, n=10 recent-onset T1D and n=2 long-standing T1D donors, belonging to three different cohorts: EUnPOD, nPOD and DiViD. Demographical/clinical characteristics of donors are reported in **Table 1**.

**Table 1.** Main demographical/clinical characteristics of donors included in the study. The following characteristics for each donor are reported: donor type (ND: non-diabetic; Aab+: autoantibody-positive; T1D-RO: type 1 diabetic recent-onset; T1D-LS: type 1 Diabetic long-standing); case ID and cohort group (EUnPOD, nPOD, DiViD); islet autoantibody-positivity; age (years); gender (F/M); Body Mass Index; diabetes duration (years or weeks from diagnosis).

| Donor Type | Case ID | Cohort | Autoantibodies | Age (years) | Gender | BMI (Kg/m<sup>2</sup>) | Diabetes Duration [years (y); weeks (w)] |
| --- | --- | --- | --- | --- | --- | --- | --- |
| ND | 311017 | EUnPOD | GADA neg, IA-2A neg, ZnT8A neg | 54 | F | 21,3 | / |
| ND | 141117 | EUnPOD | GADA neg, IA-2A neg, ZnT8A neg | 49 | M | 25,8 | / |
| ND | 181117 | EUnPOD | GADA neg, IA-2A neg, ZnT8A neg | 39 | M | 23,6 | / |
| ND | 110118 | EUnPOD | GADA neg, IA-2A neg, ZnT8A neg | 46 | F | 32,5 | / |
| ND | 210518 | EUnPOD | GADA neg, IA-2A neg, ZnT8A neg | 44 | M | 24,5 | / |
| ND | 161118 | EUnPOD | GADA neg, IA-2A neg, ZnT8A neg | 39 | F | 22,3 | / |
| ND | 110119 | EUnPOD | GADA neg, IA-2A neg, ZnT8A neg | 22 | M | 27,2 | / |
| ND | 6179 | nPOD | GADA neg, IA-2A neg, ZnT8A neg, mIAA neg | 22 | F | n/a | / |
| ND | 6178 | nPOD | GADA neg, IA-2A neg, ZnT8A neg, mIAA neg | 25 | F | n/a | / |
| ND | 6098 | nPOD | GADA neg, IA-2A neg, ZnT8A neg, mIAA neg | 18 | M | n/a | / |
| Aab+ | 6027 | nPOD | GADA neg, IA-2A neg, ZnT8A pos, mIAA neg | 19 | M | n/a | / |
| Aab+ | 6167 | nPOD | GADA neg, IA-2A pos, ZnT8A pos, mIAA neg | 37 | M | n/a | / |
| T1D-RO | Case 1 | DiViD | IA pos, GADA pos, IA-2A pos, ZnT8A pos | 25 | F | 21 | 4w |
| T1D-RO | Case 2 | DiViD | IA pos, GADA neg, IA-2A pos, ZnT8A pos | 24 | M | 20,9 | 3w |
| T1D-RO | Case 3 | DiViD | IA pos, GADA neg, IA-2A pos, ZnT8A pos | 34 | F | 23,7 | 9w |
| T1D-RO | Case 4 | DiViD | IA pos, GADA pos, IA-2A neg, ZnT8A pos | 31 | M | 25,6 | 5w |
| T1D-RO | Case 5 | DiViD | IA pos, GADA pos, IA-2A neg, ZnT8A pos | 24 | F | 28,6 | 5w |
| T1D-RO | Case 6 | DiViD | IA pos, GADA neg, IA-2A neg, ZnT8A neg | 35 | M | 26,7 | 5w |
| T1D-RO | 6113 | nPOD | GADA neg, IA-2A neg, ZnT8A neg, mIAA pos | 13 | F | n/a | 1y |
| T1D-RO | 6087 | nPOD | GADA neg, IA-2A neg, ZnT8A pos, mIAA pos | 18 | M | n/a | 4y |
| T1D-LS | 060217 | EUnPOD | GADA pos, IA-2A neg, ZnT8A neg | 39 | F | 24,5 | 21y |
| T1D-LS | 120718 | EUnPOD | GADA neg, IA-2A neg, ZnT8A neg | 81 | F | 31,2 | 45y |

#### EUnPOD cohort

Whole pancreata were obtained from brain-dead multi-organ donors within the European Network for Pancreatic Organ Donors with Diabetes (EUnPOD), in the context of the INNODIA consortium (www.innodia.eu). After acquisition of informed research consent, collected pancreata were processed using standardized procedures at University of Pisa. INNODIA EUnPOD multiorgan donors’ pancreata were obtained with approval of the local Ethics Committee at the University of Pisa. In this study we analyzed pancreatic sections from 7 non-diabetic islet cell-autoantibodies negative (ND) and 2 long-standing (LS) T1D donors.

#### nPOD cohort

Whole pancreas was obtained from brain-dead multi-organ donors within the Network for Pancreatic organ Donors with Diabetes (nPOD), founded with the support of the Juvenile Diabetes Research Foundation (JDRF, www.jdrf.npod.org). After acquisition of informed research consent, pancreas collected were processed using standardized procedures at University of Florida. In the present study we used FFPE pancreatic sections from 3 non-diabetic islet cell-autoantibody-negative (ND), 2 non diabetic islet cell-specific autoantibody positive (Aab^+^) and 2 T1D donors.

#### DiViD Cohort

Following the acquisition of appropriate consents, n=6 recent-onset (< 9 weeks from diagnosis) T1D patients underwent pancreatic biopsy by adopting laparoscopic pancreatic tail resection, in the context of the Diabetes Virus Detection (DiViD) study. The pancreatic tissue was processed for multiple purposes including FFPE tissue blocks. Collection of pancreatic tissue in the DiViD study was approved by the Norwegian Governments Regional Ethics Committee. Written informed consent was obtained from all individuals with type 1 diabetes after they had received oral and written information from the diabetologist and the surgeon separately.

### 2.2 Immunofluorescence staining

FFPE sections were analyzed using multiplexed (triple or quadruple) immunofluorescence staining followed by multicolor confocal imaging analysis and/or whole-slide scanning analysis, in order to evaluate the expression and localization of Melanoma-Differentiation-associated gene 5 (MDA5).

After deparaffinization and rehydration through a gradient ethanol series and finally distilled H_2_O, sections were incubated with Tris-Buffered saline (TBS) supplemented with 5%/donkey serum (MDA5 staining) and TBS supplemented with 5%/goat serum (insulin, glucagon, somatostatin, chromogranin A staining) to reduce non-specific reactions. Antigen retrieval was performed with 10mM Tris/1mM EDTA/0.05% Tween 20, pH 9 in boiling water for 20min. Subsequently sections were incubated with polyclonal goat anti-human MDA5 (dilution 1:100; ab4544, Abcam) over night (ON), polyclonal guinea pig anti-insulin (undiluted, IR002, Dako) for 20min, monoclonal mouse anti-glucagon (dilution 1:300, MAB1249, clone #181402, R&D Systems) or monoclonal rabbit anti-glucagon (dilution 1:100, A0565, Dako) for 1h, monoclonal rat anti-human somatostatin (dilution 1:100; MAB2358, R&D) and monoclonal rabbit anti-chromogranin (dilution 1:400, ab15160, Abcam) ON, as primary antibodies. After incubation with primary antibodies, sections were incubated with secondary antibodies, reacting with goat, mouse, guinea pig, rabbit and rat [all IgG (H+L) Alexa Fluor conjugated, dilution 1:500; Molecular Probe] for 1h or Goat anti-rabbit IgG (H+L) Brilliant Violet (dilution 1:50, Jackson ImmunoResearch) for 2h. DNA was counterstained with DAPI. Sections were finally mounted with DAKO Fluorescence Mounting Medium (S3023 Dako). Reagent details are reported in **Supplementary Table 1**.

### 2.3 Image acquisition and analysis

Images were acquired using Leica TCS SP5 confocal laser scanning microscope system (Leica Microsystems, Wetzlar, Germany) and NanoZoomer S60 Digital slide scanner (Hamamatsu Photonics K.K., Hamamatsu City, Japan). Images were analyzed using Leica Application Advance Fluorescence (LasAF) and with NDP view2 plus software. In particular, the percentage of colocalization rate of MDA5-insulin and MDA5-glucagon for each pancreatic islet was quantified determining the region of interest (ROI), drawn to calculate the “colocalization rate” (which indicates the extent of colocalization in percentage) as ratio between the colocalization area and the image foreground (in which the colocalization area represents the ratio of the area of colocalizing fluorescence signals and the image foreground represents the image area with fluorescence signal, calculated from the difference between the area of ROI and the area of image background).

### 2.4 Statistical analysis

Statistical analyses were performed using Graph Pad Prism 8 (GraphPad software, San Diego). Results were expressed as mean ± SD. Differences between groups were assessed by Mann-Whitney U test or Kruskal-Wallis multiple comparison test (for non-parametric data). We considered statistically significant a p-value less than 0,05 (p <0,05).

## 3 Results

### 3.1 Immunofluorescence analysis of pancreatic MDA5 expression in non-diabetic donors’ reveals a preferential localization in α-cells

Firstly, we sought to determine the expression and distribution of MDA5 in adult non-diabetic human pancreas using multiplex immunofluorescence. To this aim, we analyzed FFPE pancreatic tissue sections from n=10 non-diabetic/autoantibody-negative adult donors (age: 35,8 ± 12,9y; gender: 5F, 5M) belonging to EUnPOD and nPOD networks. In non-diabetic human pancreas, MDA5 was exclusively expressed in pancreatic islet endocrine cells, as shown by a near perfect colocalization between chromogranin-A and MDA5 (**Supplementary Figure 1a, panels A to L**). Rare, scattered MDA5-positive, chromogranin A-positive cells were also identified interspersed within the exocrine pancreas (**Supplementary Figure 1a, panels I to L, yellow arrow)**. In such circumstances, no MDA5-positive, chromogranin A-negative scattered cells were identified.

In order to establish the endocrine cell types expressing MDA5 within the human pancreas in non-diabetic context, we performed a triple immunofluorescence analysis aimed at detecting α- and β- cells alongside with MDA5. In human pancreatic islets, MDA5 was expressed both in α- and β-cells (**Figure 1a**, **Panels A to D and Supplementary Figure 1b**). Outside pancreatic islets, most of the scattered MDA5-positive cells were also glucagon-positive and rarely showed insulin-positivity. Moreover, no MDA5-positive, insulin- and glucagon-negative cells were identified in islets or interspersed in exocrine tissue, thus confirming that in non-diabetic human pancreas MDA5 was expressed in endocrine cells, exclusively in α- or in β-cells. Remarkably, the imaging colocalization analysis performed on a total of 147 pancreatic islets from 10 non-diabetic, autoantibody-negative adult donors, showed a significantly higher colocalization rate of MDA5-glucagon in comparison to MDA5-insulin (20,5±12,5 % vs 14,4±10,6 %, p<0,01) (**Figure 1a**, **panels A to D**, **Figure 1b, 1c**).

**Figure 1.**
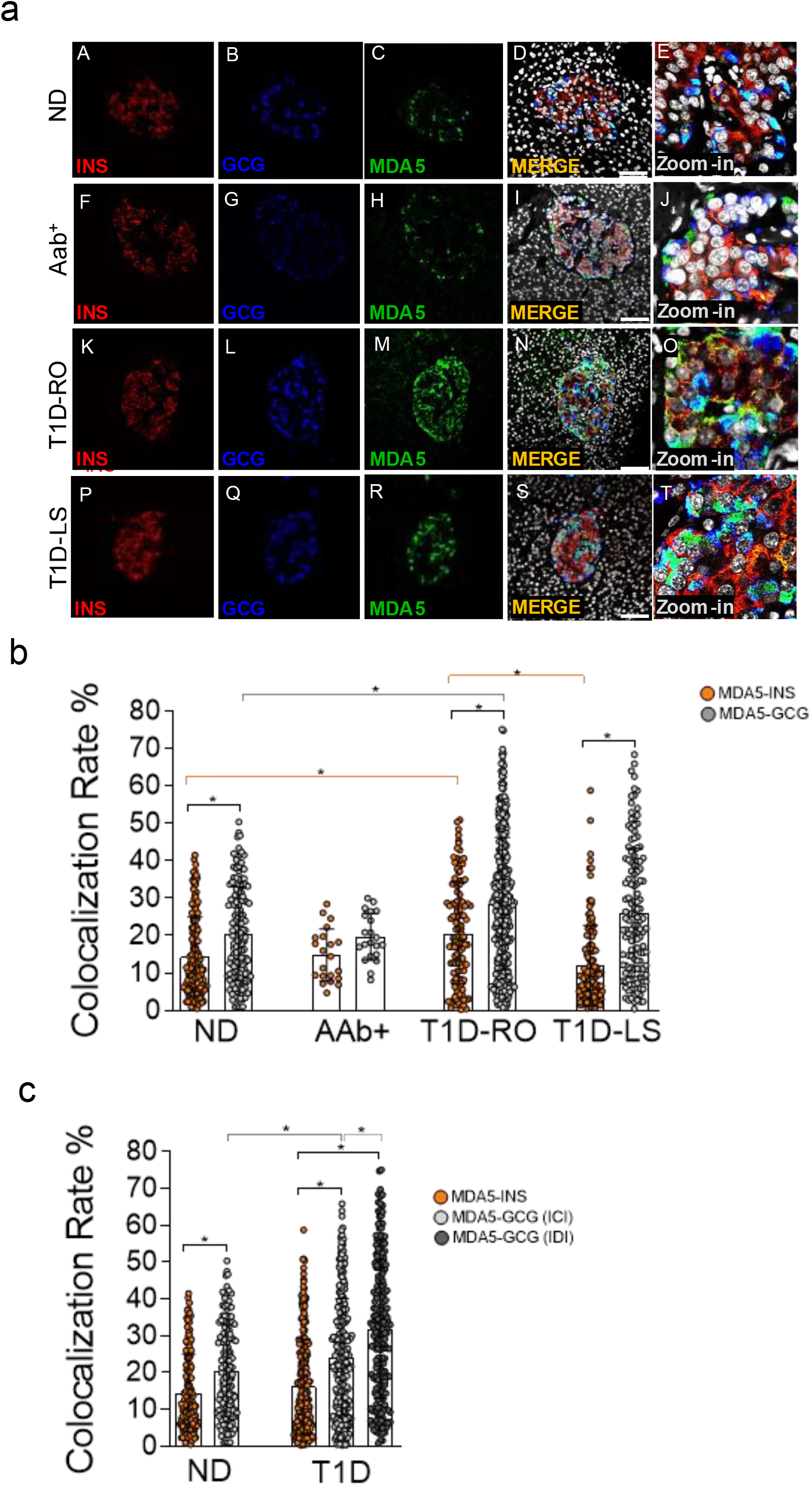
Triple Immunofluorescence analysis of insulin, glucagon and MDA5 in human pancreata of non-diabetic, Aab+, and T1D multiorgan donors. **(a)** Representative images showing fluorescence confocal microscopy imaging analysis of FFPE pancreatic tissue sections derived from non-diabetic donors (panel A-E) (n=10) (ND), autoantibody-positive donors (Panels F-J) (n=2: 6027, 6167) (Aab+), type 1 diabetic recent onset donors (Panels K-O) (T1D-RO) (n=8: DiViD cases −1, −2, −3, −4, −5, −6, nPOD 6113, 6087), and type 1 diabetic longstanding donors (panels P-T) (T1D-LS) (n=2: EUnPOD 060217, 120718), stained for insulin (red, panels A,F,K,P), glucagon (blue, panel B,G,L,Q) and MDA5 (green, panels C,H,M,R). Channels merge shows the colocalization of insulin and MDA5 in yellow/orange and of glucagon and MDA5 in turquoise color. A zoom-in inset reports details of pancreatic islets of channels merge images. Scale bars in panels D, I, N and S are 75 μm. **(b)** Colocalization rate analysis of MDA5-insulin (orange), MDA5-glucagon (light grey) in non-diabetic controls (ND) (n=10), autoantibody positive (Aab+) (n=2), T1D-RO (n=8) and T1D-LS (n=2). Each dot represents an individual pancreatic islet reported as a colocalization rate value between MDA5-glucagon or MDA5-insulin. Mean ± S.D. values are reported as a histogram plot. Statistics performed using ANOVA followed by Kruskal-Wallis Multiple Comparison test (*p<0.05). **(c)** Colocalization rate analysis of MDA5-insulin (orange), MDA5-glucagon in insulin-containing islets (ICI) (light grey), MDA5-glucagon in insulin-deficient islets (IDI) (dark grey) in non-diabetic controls (ND) (n=10) and in T1D donors (n=10, RO+LS). Each dot represents an individual pancreatic islet reported as a colocalization rate value of MDA5-glucagon or MDA5-insulin. Mean ± S.D. values are reported as a histogram plot underlying dot plot. Statistics performed using ANOVA followed by Kruskal-Wallis Multiple Comparison test (*p<0.05).

These results indicate that: (*i*) in non-diabetic human pancreas, MDA5 was expressed in endocrine cells; (*ii*) in pancreatic islets, a higher proportion of α-cells were positive for MDA5 in comparison to β-cells; (*iii*) in such context, only a small fraction (14 to 20%) of β- and α-cells were positive for MDA5.

### 3.2 MDA5 expression is increased in pancreatic islets of T1D donors

The expression and distribution of MDA5 was also analysed in pancreata obtained from autoantibody-positive (Aab^+^) and from T1D donors with different disease duration, namely: recent-onset (RO) (3 weeks to 4 years) and longstanding (LS) (>20 years) donors. Overall, MDA5 expression pattern was analysed in n=2 Aab^+^, n=8 T1D-RO (6 from DiViD study and 2 from nPOD cohort) and n=2 T1D-LS donors (from EUnPOD cohort) (**Table 1**).

In line to what previously observed in non-diabetic donors pancreata, MDA5 was expressed in α- and β-cells both in Aab^+^ and in T1D donors (**Figure 1a**). Such data were confirmed using a 3D z-scan analysis using fluorescence confocal microscopy (**Supplementary Figure 2a**), revealing the actual colocalization of MDA5 with insulin or glucagon signals, as shown by the deconvolution imaging analysis (**Supplementary Figure 2b**). Quadruple immunofluorescence analysis of chromogranin-A, insulin, glucagon and MDA5 as well as quadruple immunofluorescence analysis of somatostatin, insulin, glucagon and MDA5 in pancreatic tissue of T1D donors (**Supplementary Figure 3a and 3b**) further confirmed that MDA5-positive cells were endocrine but not somatostatin-positive, thus exclusively α- or β-cells.

The colocalization rate analysis in pancreatic islets revealed a preferential MDA5 expression in α- vs β-cells also in T1D context (27,8 ± 17,5 % vs 16,3 ± 13,0 %, n=246 ICI and n=224 IDI, p<0,0001), independently from disease duration (T1D RO: 28,5±17% vs 20,6±13,5%; T1D-LS: 26,1±17,1% vs 11,9±10,8%). Such pattern occurred in Aab^+^ donors as well.

Importantly, pancreatic islets of T1D-RO donors showed a higher colocalization rate MDA5-glucagon as well as MDA5-insulin in comparison to non-diabetic controls (**Figure 1b**)(28,5±17% vs 20,5±12,5 % and 20,6±13,5%; vs 14,4±10,6 %, respectively; p< 0.05). Such increase was more pronounced in β-cells of T1D-RO respect to those of T1D-LS donors who showed a similar MDA5-insulin colocalization rate respect to non-diabetic controls (**Figure 1b**).

As expected, a case by case analysis of MDA5-glucagon, MDA5-insulin colocalization rate values highlighted the elevated heterogeneity among islets and among different donors (**Supplementary Figure 4**), even though the increased proportion of α- and β-cells positive for MDA5 in T1D cases vs non-diabetic controls was consistent. Of note, DiViD case-3 showed the highest MDA5-insulin colocalization rate among DiViD cases (**Supplementary Figure 4**).

Interestingly, an additional analysis of MDA5-glucagon and MDA5-insulin colocalization rate in T1D-RO and T1D-LS performed taking into consideration ICIs and IDIs separately, revealed that in IDIs colocalization rate MDA5-glucagon was higher respect to ICIs **(Figure 1c**).

### 3.3 Identification of MDA5-positive/hormones-negative islet-like structures in pancreas of recent-onset T1D donors

In all T1D-RO donors [DiViD (n=6) and nPOD (n=2)], we observed several cells showing MDA5-positive signal, though negative for both insulin and glucagon; on the contrary, in non-diabetic control donors pancreas, we did not observe MDA5-positive, insulin- and glucagon-negative cells, neither within the islets parenchyma nor scattered in the exocrine. (**Figure 2a**).

**Figure 2.**
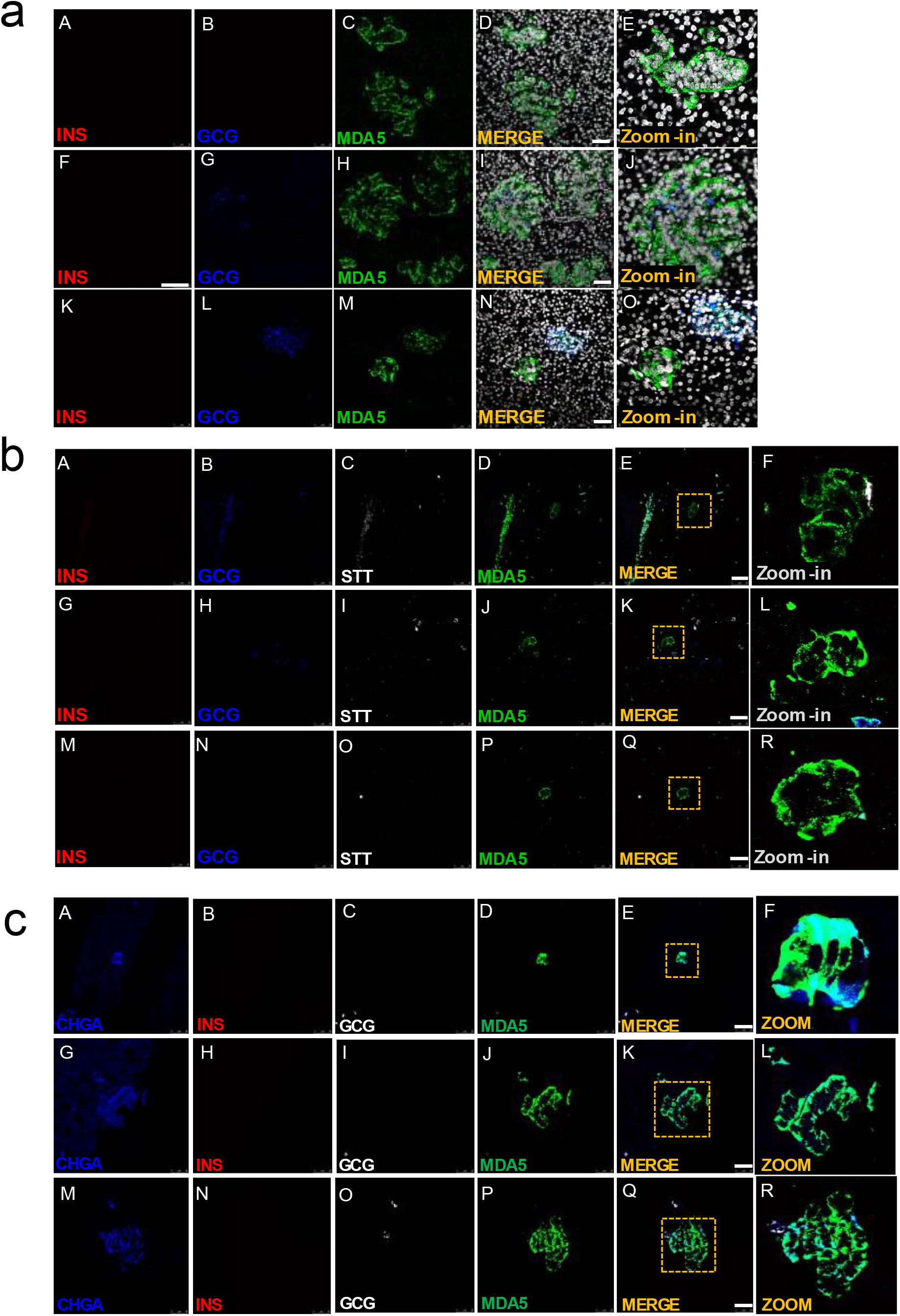
Immunofluorescence detection of MDA5-positive/hormones-negative islet-like structures in pancreas of T1D donors. **(a)** Representative images reporting triple immunofluorescence analysis of insulin (red, panels A, F, K), glucagon (blue, panels B, G, L), MDA5 (green, panels C, H, M), DAPI (white) in T1D-RO donors (nPOD 6113: panels A-E; nPOD 6087: panels F-J; DiViD case 3: panels K-O. Zoom-in insets (panels E, J, O) report details of channels merge images. Scale bars in panels D, I, N are 50 μm. **(b)** Representative images of quadruple immunofluorescence analysis of insulin (red, panels A, G, M), glucagon (blue, panels B, H, N), SST (white, panels C, I, O) and MDA5 (green, panels D, J, P) in T1D-RO donors. Zoom-in insets (panels F, L, R) report details of channels merge images. Scale bars in panels E, K, Q are 50 μm. **(c)** Representative images of quadruple immunofluorescence analysis of chromogranin A (blue, panels A, G, M), insulin (red, panels B, H, N), glucagon (white, panels C, I, O) and MDA5 (green, panels D, J, P) in T1D-RO donors. Zoom-in insets (panels F, L, R) report details of channels merge images. Scale bars panels E, K, Q = 50 μm.

Interestingly, such cells were histologically organized in structures reminiscent of the islet architecture (**Figure 2a**, **panel E**), whose diameter and cell numbers varied consistently (**Figure 2a**, **panels I, D, N**). In some we observed several glucagon-positive cells, surrounded by a consistent number of MDA5-positive, insulin- and glucagon negative cells (**Figure 2a**, **panels L, J),** potentially suggesting an islet lineage.

A Z-scan confocal imaging analysis further confirmed that such structures were MDA5-positive but insulin and glucagon-negative, also upon 3D scanning imaging and a deconvolution projection analysis (**Supplementary Figure 5a and 5b**).

Strikingly, MDA5-positive cells belonging to such structures were somatostatin-negative (**Figure 2b**, **panels A-R**) but clearly positive for chromogranin A, thus definitely suggesting their endocrine origin (**Figure 2c**, **panels A-R**). In addition, in such structures, the MDA5 fluorescence signal was visibly higher than what previously observed for α- or β-cells, thus suggesting an elevated expression of MDA5 in these cells.

Finally, a detailed observation of pancreatic tissue sections stained for insulin, glucagon and MDA5 revealed that such structures were present exclusively in T1D-RO donors (both DiViD and nPOD) but absent in T1D-LS (EUnPOD) or in non-diabetic control donors (both EUnPOD and nPOD).

### 3.4 Distribution analysis of MDA5-positive islets in pancreas of T1D DiViD donors

In order to evaluate the distribution and eventual heterogeneity of pancreatic islets in terms of MDA5 expression, we performed a whole slide scanning imaging analysis of pancreatic tissue sections derived from EUnPOD non-diabetic donors and from T1D DiViD cases, stained for insulin, glucagon and MDA5 (**Figure 3a**, **panel A**). Whole slide fluorescence scanning imaging analysis has the advantage to allow the exploration of the entire pancreatic tissue section.

**Figure 3.**
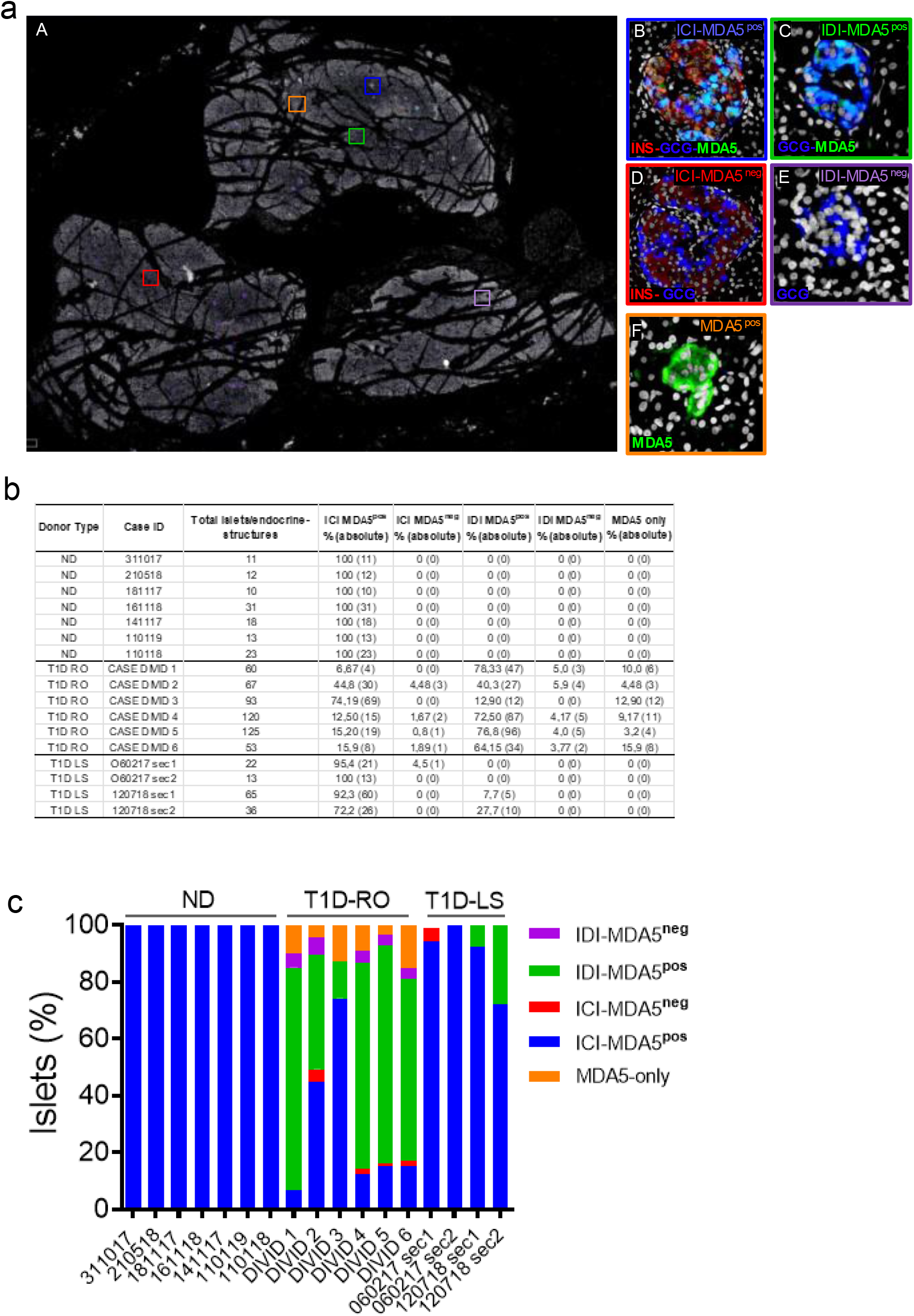
Whole-slide imaging islets distribution analysis based on MDA5 positivity in ND and T1D donors. **(a)** Representative whole slide image of T1D case-3 of a triple immunofluorescence for insulin (red), glucagon (blue), MDA5 (green) and DAPI (white) analyzed using Nanozoomer S60 slide scanner. The image (panel A) reports the location of 5 different histological structures identified in T1D pancreatic sections. Zoom-in insets for each type of islet or islet-like structure are reported on the left. Blue square (panel B): Insulin-containing islet (ICI), MDA5-positive; red square (panel D): Insulin-containing islet, MDA5-negative; green square (panel C): Insulin-deficient islets, MDA5-positive; magenta square (panel E): Insulin-deficient islets, MDA5-negative; orange square: MDA5-only positive islet-like structures. **(b)** Table showing islet distribution based on MDA5-positivity in pancreatic tissue sections of ND, T1D-RO and T1D-LS. Results are reported as percentage values (alongside with absolute values) over the total number of islets detected. Donor types as well as cases IDs are reported. **(c)** Graphical representation of the distribution of pancreatic islets based on MDA5 positivity in pancreatic tissue sections of ND, T1D-RO (DiViD cases) and T1D-LS.

In non-diabetic donors, ICIs were all positive for MDA5. No MDA5-positive/hormones-negative islet-like structures were identified in non-diabetic context, confirming what previously observed by confocal microscopy.

In T1D DiViD donors, we found a high heterogeneity among pancreatic islets, including the presence of ICIs or IDIs-MDA5 negative (**Figure 3a**, **panels C-D**). Overall, in T1D DiViD cases a large fraction of pancreatic islets were positive for MDA5, and only a minor proportion of them showed very low or null expression of MDA5 (**Figure 3b-c**). In DiViD cases, a variable percentage of MDA5-chromogranin A positive/hormones-negative islet-like structures (over the total number of islets detected) were identified, ranging from 3,2% (DiViD case-5) to 15,9% (DiViD case-6). (**Figure 3b**).

Collectively, these results suggest that: (*i*) in T1D donors, human pancreatic islets show a higher proportion of α- and β-cells expressing MDA5 respect to non-diabetic controls or T1D-LS donors; (*ii*) in T1D, both α- and β-cells drive the increase of MDA5 positive cells in human pancreatic islets; (*iii*) such increase is more pronounced in T1D-RO respect to T1D-LS; (iv) in T1D-RO donors, we uncovered the existence of MDA5-expressing islet-like structures which were positive for chromogranin A but negative for islet main hormones (insulin, glucagon and somatostatin).

## 4 Discussion

The hypothesis that viruses could play a key role in the pathogenesis of T1D has a long history. Enteroviruses (EVs), especially Coxsackievirus B-group (CVB), have been associated to the initiation and/or acceleration of pancreatic islet autoimmunity (24). As a matter of fact, many evidence have shown the ability of EVs to infect human pancreatic β-cells and to impair their function and/or survival (4–8). Pancreatic immunohistochemical data showed the selective positivity of Viral Capsid Protein-1 (VP1) in β-cells. Such selectivity is corroborated by additional evidence showing: (*i*) a strong permissiveness of β-cells to EVs when infected in-vitro (25,26), and (*ii*) the specific expression of Coxsackie and Adenovirus receptor (CAR) in β-cells (12). Of note, although a strong causal association between EV infections and T1D is still debated, several mechanisms have been proposed, including: (i) direct destruction of β-cells, (ii) by-stander activation of autoreactive T cells, (iii) molecular mimicry, and viral persistence causing β-cell dysfunction.

A crucial piece of the puzzle is represented by intracellular antiviral molecules involved in innate immune response, which characterize the first line of defense by coordinating signaling pathways activation leading to inflammatory responses. Among others, MDA5 has been identified as an important component of the intracellular innate immune response to EV intermediate dsRNAs, which is necessary to prevent early replication of the virus and to limit tissue damage though a rapid and efficient pathogen clearance (21).

Although MDA5 expression and distribution have been evaluated in murine pancreas and in pancreatic samples of multiorgan donors affected by fulminant diabetes (27), a detailed characterization of MDA5 expression in human T1D and non-diabetic pancreas is still missing. We took advantage from several organ biorepository networks in order to analyze the MDA5 expression pattern in pancreatic tissues from T1D donors with different disease duration, as well as from non-diabetic/autoantibody-positive and autoantibody-negative donors. This has been accomplished by accessing three different biorepositories: EUnPOD, nPOD and DiViD study biobank.

Firstly, we analyzed the expression pattern of MDA5 in pancreas of control donors. Our results showed that, in non-diabetic pancreas, MDA5 was specifically expressed in endocrine cells, particularly in α- and in β-cells, thus confirming that islet cells are equipped to efficiently respond to viral RNA intermediates. Of interest, in non-diabetic context, a higher proportion of α-cells expressed MDA5 respect to β-cells. Such results are concordant with different bulk RNA transcriptomics datasets of sorted human α- and β-cells (22,28,29) showing a preferential expression of MDA5 mRNA in α-cells respect to β-cells; additionally, these data confirm the bona-fide of the MDA5 antibody used in the present study.

The colocalization rate of MDA5-glucagon and MDA5-insulin in non-diabetic controls was highly heterogenous both among islets of the same donor and among different donors; nevertheless the overall result showed a significant preferential localization in α-cells. Such heterogeneity was expected, based on previous findings on other molecules (30,31) and on other reports (32,33).

The preferential localization of MDA5 in α-cells suggests that this cell type is better equipped than β-cells to efficiently respond to EVs entry by promptly activating pro-inflammatory signaling pathways which contribute to virus clearance in the early stages of infection. Such observation is substantiated by in-vitro studies showing a higher capacity of α-cells vs β-cells to increase the expression of MDA5 and of other key antiviral proteins, namely STAT1 and MX1, in response to CVB infection (21). As a matter of fact, MDA5 has been found to play a key role in IFNs-I- and IFNs-III induction and in inflammatory cytokines production following virus infection; consequently, synthesis of pro-inflammatory mediators may lead to the expression of ISGs, whose products direct antiviral and immunoregulatory effects.. On the contrary, lack of a fully functional antiviral signaling machinery characterized by MDA5 loss of function, absence or reduced expression could be linked to the establishment of a subtle chronic inflammatory response and to a long-lasting non-cytolytic infection which may activate autoimmunity.

Our results also showed a higher proportion of α- and β-cells expressing MDA5 in recent onset T1D cases respect to non-diabetic controls. The increased colocalization rate between MDA5-glucagon and MDA5-insulin in T1D cases, possibly reflects ongoing inflammation of pancreatic islets particularly in recent onset cases. Of note, the strongest MDA5 increase was observed in β-cells of T1D DiViD case-3 which, among T1D DiViD donors, previously showed high level of insulitis alongside with a strong expression of HLA class-I molecules and several specific cytokines and chemokines (30,34,35). These results are in line with in-vitro studies showing that increased expression of MDA5 is also driven by pro-inflammatory molecules. Of note, it has been demonstrated that exposure of pancreatic islet cells to viruses or to viral intermediates (dsRNAs) leads to MDA5 increased expression (16,20); remarkably, evidence of EVs infection has been detected in pancreatic islets of all DiViD cases across different laboratories employing multiple approaches (6,10).

An additional result worth of mention is the higher colocalization rate MDA5-glucagon in IDI vs ICI. Based on the fact that IDIs are deprived of β-cells, thus less inflamed than ICIs, we can hypothesize that the increase of MDA5 in IDIs could be due to other factors than pro-inflammatory molecules activity. It has been reported that in specific conditions MDA5 could also sense endogenous dsRNAs generated by epigenetic activation of repeated genomic DNA sequences. Indeed, increased MDA5 expression and activation has been reported in cancers treated with de-methylating agents, leading to the accumulation of RNAs (36,37). Physiologically, this phenomenon could be due to phenotypic changes requiring activation of epigenetic mechanisms and RNA editing modifications. As such, we can hypothesize that residual α-cells within IDIs are subjected to strong epigenetic pressure with the aim to modify their phenotype, thus leading to the transcription of dsRNAs or to modifications of RNA editing mechanisms which finally increase MDA5 expression and activity. Importantly, it has been recently shown that rodent pancreatic α-cells undergo massive transcriptional changes upon total β-cells ablation, including upregulation of genes involved in interferon signaling and proliferation (38); of note, an increased expression of MDA5 as early as 5 days post β-cells ablation (38)(see GEO dataset GSE155519).

Strikingly, for the first time, we also detected several islet-like cell clusters which were positive for MDA5 but negative for the main islet hormones (insulin, glucagon and somatostanin). Such cells were of endocrine origin since they showed positivity for chromogranin A. Although we did not test for other minor hormones (pancreatic-polypeptide, ghrelin), the elevated number of cells composing these islet-like clusters allowed us to exclude that they could be δ- or ε-cells. In addition, given their morphology, structure, nuclei conformation and, more importantly, the presently shown positivity for Chromogranin-A, we can exclude their immune origin.

Increased frequency of chromogranin A-positive/hormones-negative cells has been documented in pancreas of rodent models with diabetes, in donors with T1D and T2D (39–41), as well as in human pancreatic tissues derived from donors affected by pancreatitis (42). Here we showed that such chromogranin A-positive/ hormone-negative cell clusters were also strongly positive for MDA5. The exact origin of these islet-like clusters has yet to be elucidated. In line to what previously suggested, we can hypothesize that these clusters could be derived from pancreatic islets cells undergoing to dedifferentiation phoenomena. Interestingly, it has been shown that human β-cells dedifferentiate upon virus-like infection simulated in-vitro by PolyI:C (PIC) treatment (43); of note, upon PIC exposure, it an increased MDA5 expression alongside with a reduced expression of several genes related to β-cells identity and function was observed (43). Similarly, it has been shown that pro-inflammatory molecules may induce β-cells dysfunction and phenotype loss, together with an induction of MDA5 expression. Among pro-inflammatory molecules tested, IL-1β was shown to be the most potent inducer of in-vitro dedifferentiation (44). In support of a role of inflammation in promoting islet dedifferentiation and appearance of MDA5-positive/hormones-negative endocrine cells, we observed that one of the highest frequency of MDA5-positive/hormone-negative islet-like clusters was detected in pancreas of T1D DiViD case-3 which also showed high rate of inflammation, thus corroborating the current view of an inflammation-driven mechanism of dedifferentiation induction. Strikingly, MDA5-positive/hormone-negative islet-like clusters were not observed in autoantibody-positive non-diabetic donors and in long-standing T1D donors, usually characterized by lower levels of inflammation respect to recent-onset donors.

Alternatively, we can speculate that MDA5-positive/hormones-negative endocrine cells may represent newly forming islets, recapitulating what observed in human neonatal pancreas enriched in chromogranin-A positive/hormone-negative structures.

Whatever the origin of these islet-like clusters, a function for MDA5 in putatively undifferentiated cells should be deciphered. It is possible that high MDA5 expression found in these clusters may be the consequence of inflammatory phoenomena leading to activation of specific innate signaling pathways and, in parallel, causing dedifferentiation. On the other hand, MDA5 increased expression and activity could also be due to epigenetic changes occurring during phenotypic modifications taking place throughout dedifferentiation, causing RNAs accumulation and sensing by MDA5.

In conclusion, we showed that MDA5 is expressed in pancreatic endocrine cells with a preferential localization in α-cells vs β-cells, remarkably suggesting that α-cells are better equipped respect to β-cells to respond and activate viral clearance mechanism; such difference may render β-cells more susceptible to the establishment of persistent low-grade viral infections. We also showed that MDA5 is increased in recent onset T1D donors possibly in consequence of elevated inflammatory conditions. Of note, in these donors we highlighted the identification of MDA5-positive/hormones-negative endocrine cell clusters thus opening to novel roles for MDA5 in putative dedifferentiation mechanisms in T1D.

## 6 Author Contributions

*LN, NB, and GS performed the experiments, analyzed the data, and wrote the manuscript. GG, DF and GL analyzed the data and contributed to the scientific discussion. CF contributed to the scientific discussion. LN, GS, and FD reviewed the manuscript and designed experiments. LK and KD provided support for DiViD cohort and contributed to the scientific discussion. LM and PM reviewed the manuscript, provided support for EUnPOD donors, and contributed to the scientific discussion. All authors contributed to the article and approved the submitted version*

## 8 Acknowledgments

The secretarial help of Maddalena Prencipe and Alessandra Mechini was highly appreciated

## Data Availability Statement

All datasets generated for this study are included in the article/SupplementaryMaterial

## Reference

1. Atkinson MA, Eisenbarth GS, Michels AW. Type 1 diabetes. Lancet. 2014 Jan 4;383(9911):69–82.

2. Spagnuolo I, Patti A, Sebastiani G, Nigi L, Dotta F. The case for virus-induced type 1 diabetes. Curr Opin Endocrinol Diabetes Obes. 2013 Aug;20(4):292–298.

3. Galleri L, Sebastiani G, Vendrame F, Grieco FA, Spagnuolo I, Dotta F. Viral infections and diabetes. Adv Exp Med Biol. 2012;771:252–271.

4. Dotta F, Censini S, van Halteren AGS, Marselli L, Masini M, Dionisi S, et al. Coxsackie B4 virus infection of beta cells and natural killer cell insulitis in recent-onset type 1 diabetic patients. Proc Natl Acad Sci USA. 2007 Mar 20;104(12):5115–5120.

5. Richardson SJ, Willcox A, Bone AJ, Foulis AK, Morgan NG. The prevalence of enteroviral capsid protein vp1 immunostaining in pancreatic islets in human type 1 diabetes. Diabetologia. 2009 Jun;52(6):1143–1151.

6. Krogvold L, Edwin B, Buanes T, Frisk G, Skog O, Anagandula M, et al. Detection of a low-grade enteroviral infection in the islets of langerhans of living patients newly diagnosed with type 1 diabetes. Diabetes. 2015 May;64(5):1682–1687.

7. Richardson SJ, Leete P, Bone AJ, Foulis AK, Morgan NG. Expression of the enteroviral capsid protein VP1 in the islet cells of patients with type 1 diabetes is associated with induction of protein kinase R and downregulation of Mcl-1. Diabetologia. 2013 Jan;56(1):185–193.

8. Willcox A, Richardson SJ, Bone AJ, Foulis AK, Morgan NG. Immunohistochemical analysis of the relationship between islet cell proliferation and the production of the enteroviral capsid protein, VP1, in the islets of patients with recent-onset type 1 diabetes. Diabetologia. 2011 Sep;54(9):2417–2420.

9. Geravandi S, Richardson S, Pugliese A, Maedler K. Localization of enteroviral RNA within the pancreas in donors with T1D and T1D-associated autoantibodies. Cell Reports Medicine. 2021;

10. Oikarinen S, Krogvold L, Edwin B, Buanes T, Korsgren O, Laiho JE, et al. Characterisation of enterovirus RNA detected in the pancreas and other specimens of live patients with newly diagnosed type 1 diabetes in the DiViD study. Diabetologia. 2021 Aug 14;

11. Marroqui L, Perez-Serna AA, Babiloni-Chust I, Dos Santos RS. Type I interferons as key players in pancreatic β-cell dysfunction in type 1 diabetes. Int Rev Cell Mol Biol. 2021 Mar 23;359:1–80.

12. Ifie E, Russell MA, Dhayal S, Leete P, Sebastiani G, Nigi L, et al. Unexpected subcellular distribution of a specific isoform of the Coxsackie and adenovirus receptor, CAR-SIV, in human pancreatic beta cells. Diabetologia. 2018 Aug 3;61(11):2344–2355.

13. Apaolaza PS, Balcacean D, Zapardiel-Gonzalo J, Nelson G, Lenchik N, Akhbari P, et al. Islet expression of type I interferon response sensors is associated with immune infiltration and viral infection in type 1 diabetes. Sci Adv. 2021 Feb 24;7(9).

14. Aida K, Nishida Y, Tanaka S, Maruyama T, Shimada A, Awata T, et al. RIG-I- and MDA5-initiated innate immunity linked with adaptive immunity accelerates beta-cell death in fulminant type 1 diabetes. Diabetes. 2011 Mar;60(3):884–889.

15. Dias Junior AG, Sampaio NG, Rehwinkel J. A balancing act: MDA5 in antiviral immunity and autoinflammation. Trends Microbiol. 2019 Jan;27(1):75–85.

16. Colli ML, Moore F, Gurzov EN, Ortis F, Eizirik DL. MDA5 and PTPN2, two candidate genes for type 1 diabetes, modify pancreatic beta-cell responses to the viral by-product double-stranded RNA. Hum Mol Genet. 2010 Jan 1;19(1):135–146.

17. Nejentsev S, Walker N, Riches D, Egholm M, Todd JA. Rare variants of IFIH1, a gene implicated in antiviral responses, protect against type 1 diabetes. Science. 2009 Apr 17;324(5925):387–389.

18. Smyth DJ, Cooper JD, Bailey R, Field S, Burren O, Smink LJ, et al. A genome-wide association study of nonsynonymous SNPs identifies a type 1 diabetes locus in the interferon-induced helicase (IFIH1) region. Nat Genet. 2006 Jun;38(6):617–619.

19. Concannon P, Onengut-Gumuscu S, Todd JA, Smyth DJ, Pociot F, Bergholdt R, et al. A human type 1 diabetes susceptibility locus maps to chromosome 21q22.3. Diabetes. 2008 Oct;57(10):2858–2861.

20. Domsgen E, Lind K, Kong L, Hühn MH, Rasool O, van Kuppeveld F, et al. An IFIH1 gene polymorphism associated with risk for autoimmunity regulates canonical antiviral defence pathways in Coxsackievirus infected human pancreatic islets. Sci Rep. 2016 Dec 21;6:39378.

21. Marroqui L, Lopes M, dos Santos RS, Grieco FA, Roivainen M, Richardson SJ, et al. Differential cell autonomous responses determine the outcome of coxsackievirus infections in murine pancreatic α and β cells. Elife. 2015 Jun 10;4:e06990.

22. Eizirik DL, Sammeth M, Bouckenooghe T, Bottu G, Sisino G, Igoillo-Esteve M, et al. The human pancreatic islet transcriptome: expression of candidate genes for type 1 diabetes and the impact of pro-inflammatory cytokines. PLoS Genet. 2012 Mar 8;8(3):e1002552.

23. Lincez PJ, Shanina I, Horwitz MS. Reduced expression of the MDA5 Gene IFIH1 prevents autoimmune diabetes. Diabetes. 2015 Jun;64(6):2184–2193.

24. Dotta F, Sebastiani G. Enteroviral infections and development of type 1 diabetes: The Brothers Karamazov within the CVBs. Diabetes. 2014 Feb;63(2):384–386.

25. Ylipaasto P, Klingel K, Lindberg AM, Otonkoski T, Kandolf R, Hovi T, et al. Enterovirus infection in human pancreatic islet cells, islet tropism in vivo and receptor involvement in cultured islet beta cells. Diabetologia. 2004 Feb;47(2):225–239.

26. Frisk G, Diderholm H. Tissue culture of isolated human pancreatic islets infected with different strains of coxsackievirus B4: assessment of virus replication and effects on islet morphology and insulin release. Int J Exp Diabetes Res. 2000;1(3):165–175.

27. Aida K, Nishida Y, Tanaka S, Maruyama T. RIG-I–and MDA5-initiated innate immunity linked with adaptive immunity accelerates β-cell death in fulminant type 1 diabetes. Diabetes. 2011;

28. Dorrell C, Schug J, Lin CF, Canaday PS, Fox AJ, Smirnova O, et al. Transcriptomes of the major human pancreatic cell types. Diabetologia. 2011 Nov;54(11):2832–2844.

29. Bramswig NC, Everett LJ, Schug J, Dorrell C, Liu C, Luo Y, et al. Epigenomic plasticity enables human pancreatic α to β cell reprogramming. J Clin Invest. 2013 Mar;123(3):1275–1284.

30. Nigi L, Brusco N, Grieco GE, Licata G, Krogvold L, Marselli L, et al. Pancreatic Alpha-Cells Contribute Together With Beta-Cells to CXCL10 Expression in Type 1 Diabetes. Front Endocrinol (Lausanne). 2020 Sep 15;11:630.

31. Fignani D, Licata G, Brusco N, Nigi L, Grieco GE, Marselli L, et al. SARS-CoV-2 Receptor Angiotensin I-Converting Enzyme Type 2 (ACE2) Is Expressed in Human Pancreatic β-Cells and in the Human Pancreas Microvasculature. Front Endocrinol (Lausanne). 2020 Nov 13;11:596898.

32. Arrojo E Drigo R, Roy B, MacDonald PE. Molecular and functional profiling of human islets: from heterogeneity to human phenotypes. Diabetologia. 2020 Oct;63(10):2095–2101.

33. Damond N, Engler S, Zanotelli VRT, Schapiro D, Wasserfall CH, Kusmartseva I, et al. A map of human type 1 diabetes progression by imaging mass cytometry. Cell Metab. 2019 Mar 5;29(3):755–768.e5.

34. Richardson SJ, Rodriguez-Calvo T, Gerling IC, Mathews CE, Kaddis JS, Russell MA, et al. Islet cell hyperexpression of HLA class I antigens: a defining feature in type 1 diabetes. Diabetologia. 2016 Nov;59(11):2448–2458.

35. Reddy S, Krogvold L, Martin C, Holland R, Choi J, Woo H, et al. Distribution of IL-1β immunoreactive cells in pancreatic biopsies from living volunteers with new-onset type 1 diabetes: comparison with donors without diabetes and with longer duration of disease. Diabetologia. 2018 Mar 27;61(6):1–12.

36. Roulois D, Loo Yau H, Singhania R, Wang Y, Danesh A, Shen SY, et al. DNA-Demethylating Agents Target Colorectal Cancer Cells by Inducing Viral Mimicry by Endogenous Transcripts. Cell. 2015 Aug 27;162(5):961–973.

37. Chiappinelli KB, Strissel PL, Desrichard A, Li H, Henke C, Akman B, et al. Inhibiting DNA Methylation Causes an Interferon Response in Cancer via dsRNA Including Endogenous Retroviruses. Cell. 2015 Aug 27;162(5):974–986.

38. Oropeza D, Cigliola V, Romero A, Chera S, Rodríguez-Seguí SA, Herrera PL. Stage-specific transcriptomic changes in pancreatic α-cells after massive β-cell loss. BMC Genomics. 2021 Aug 2;22(1):585.

39. Saleh Md Moin A, Dhawan S, Cory M, Butler PC, Rizza RA, Butler AE. Increased frequency of hormone negative and polyhormonal endocrine cells in lean individuals with type 2 diabetes. J Clin Endocrinol Metab. 2016 Jul 29;101(10):jc20162496.

40. Md Moin AS, Dhawan S, Shieh C, Butler PC, Cory M, Butler AE. Increased Hormone-Negative Endocrine Cells in the Pancreas in Type 1 Diabetes. J Clin Endocrinol Metab. 2016 Jun 14;101(9):3487–3496.

41. Md Moin AS, Cory M, Ong A, Choi J, Dhawan S, Butler PC, et al. Pancreatic nonhormone expressing endocrine cells in children with type 1 diabetes. Journal of the Endocrine Society. 2017 May 1;1(5):385–395.

42. Moin ASM, Cory M, Choi J, Ong A, Dhawan S, Dry SM, et al. Increased Chromogranin A-Positive Hormone-Negative Cells in Chronic Pancreatitis. J Clin Endocrinol Metab. 2018 Jun 1;103(6):2126–2135.

43. Oshima M, Knoch K-P, Diedisheim M, Petzold A, Cattan P, Bugliani M, et al. Virus-like infection induces human β cell dedifferentiation. JCI Insight. 2018 Feb 8;3(3).

44. Nordmann TM, Dror E, Schulze F, Traub S, Berishvili E, Barbieux C, et al. The Role of Inflammation in β-cell Dedifferentiation. Sci Rep. 2017 Jul 24;7(1):6285.

