## Supplementary material for "Increased expression of viral sensor MDA5 in pancreatic islets and in hormone-negative endocrine cells in recent onset type 1 diabetic donors"

### Supplementary Table 1. Resources Table.

### Table reporting methodological details and commercial information of primary, secondary antibodies and additional reagents used in the present study

| **Reagent** | **Clonality** | **Type** | **Company** | **Cat No** | **Clone** | **Species** | **Conjugate** | **Reactivity** | **[Stock]** | **Dilution** | **Incubation time** | **Incubation Buffer** |
| --- | --- | --- | --- | --- | --- | --- | --- | --- | --- | --- | --- | --- |
| MDA5 | Polyclonal | Primary | Abcam | Ab4544 | n/a | Goat | n/a | Human | 0,5 mg/ml | 1:100 | ON | TBS/5% Donkey serum albumin |
| Polyclonal Insulin | Polyclonal | Primary | Dako | IR002 | n/a | Guinea Pig | n/a | Human, mouse, rat | n/a | undiluted | 20 min | n/a |
| Glucagon | Monoclonal | Primary | R&D System | MAB1249 | 181402 | Mouse | n/a | Human, mouse | n/a | 1:300 | 1h | TBS/5% Goat serum albumin |
| Glucagon | Polyclonal | Primary | Dako | A0565 | n/a | Rabbit | n/a | Human, mouse, rat | 35,2 g/L | 1:100 | 1h | TBS/5% Goat serum albumin |
| Somatostatin | Monoclonal | Primary | R&D System | MAB2354 | 906552 | Rat | n/a | Human, mouse | 0,1 mg/ml | 1:100 | 1h | TBS/5% Goat serum albumin |
| Chromogranin | Polyclonal | Primary | Abcam | Ab15160 | n/a | Rabbit | n/a | Human | 0,2 mg/ml | 1:400 | ON | TBS/5% Donkey serum albumin |
| Donkey ɑ-Goat IgG (H+L) | Polyclonal | Secondary | Life Technologies | A11055 | n/a | Donkey | Alexa Fluor ® 488 | Goat | 2 mg/ml | 1:500 | 1h | TBS |
| Goat ɑ-guinea-pig IgG (H+L) | Polyclonal | Secondary | Life Technologies | A21435 | n/a | Goat | Alexa Fluor ® 555 | Guinea pig | 2 mg/ml | 1:500 | 1h | TBS |
| Goat ɑ-rat IgG (H+L) | Polyclonal | Secondary | Life Technologies | A21247 | n/a | Goat | Alexa Fluor ® 647 | Rat | 2 mg/ml | 1:500 | 1h | TBS |
| Goat ɑ-mouse IgG (H+L) | Polyclonal | Secondary | Life Technologies | A21236 | n/a | Goat | Alexa Fluor ® 647 | Mouse | 2 mg/ml | 1:500 | 1h | TBS |
| Goat ɑ-rabbit IgG (H+L) | Polyclonal | Secondary | Jackson ImmunoResearch | 111-675-144 | n/a | Goat | Brilliant Violet 421 | Rabbit | 1mg/ml | 1:50 | 2h | TBS |
| 4′,6-Diamidino-2-phenylindole dihydrochloride (DAPI) | n/a | n/a | Sigma Aldrich | D8517 | n/a | n/a | n/a | n/a | 1mg/ml | 1:3000 | 5min | TBS |

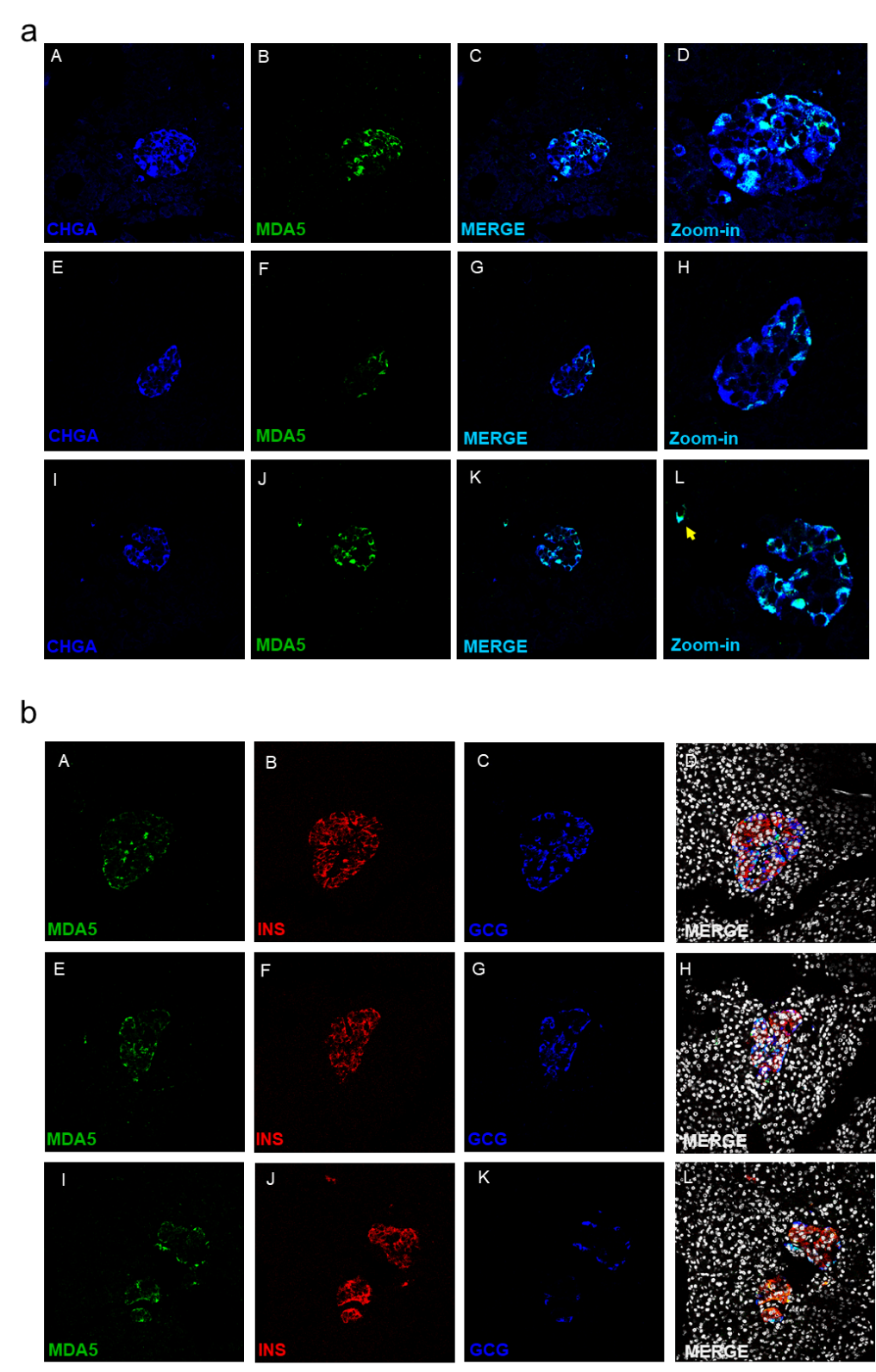

**Supplementary Figure 1. a)** Representative images of pancreatic islets within tissue sections of non-diabetic donors immunostained to detect chromogranin A (blue, panels A, E, I) and MDA5 (green, panels B, F, J). Merge panels (panels C, G, K) and Zoom-in images (panels D, H, L) show the colocalization between MDA5 and chromogranin A in light blue. The yellow arrow indicate a chromogranin A-positive/MDA5-positive cell scattered in the exocrine tissue. Three different pancreatic islets belonging to different donors are reported.

**b**) Representative images of pancreatic islets within tissue sections of non-diabetic donors immunostained to detect MDA5 (green, panels A, E, I), insulin (red, Panels B, F, J) and glucagon (blue, panels C, G, K).

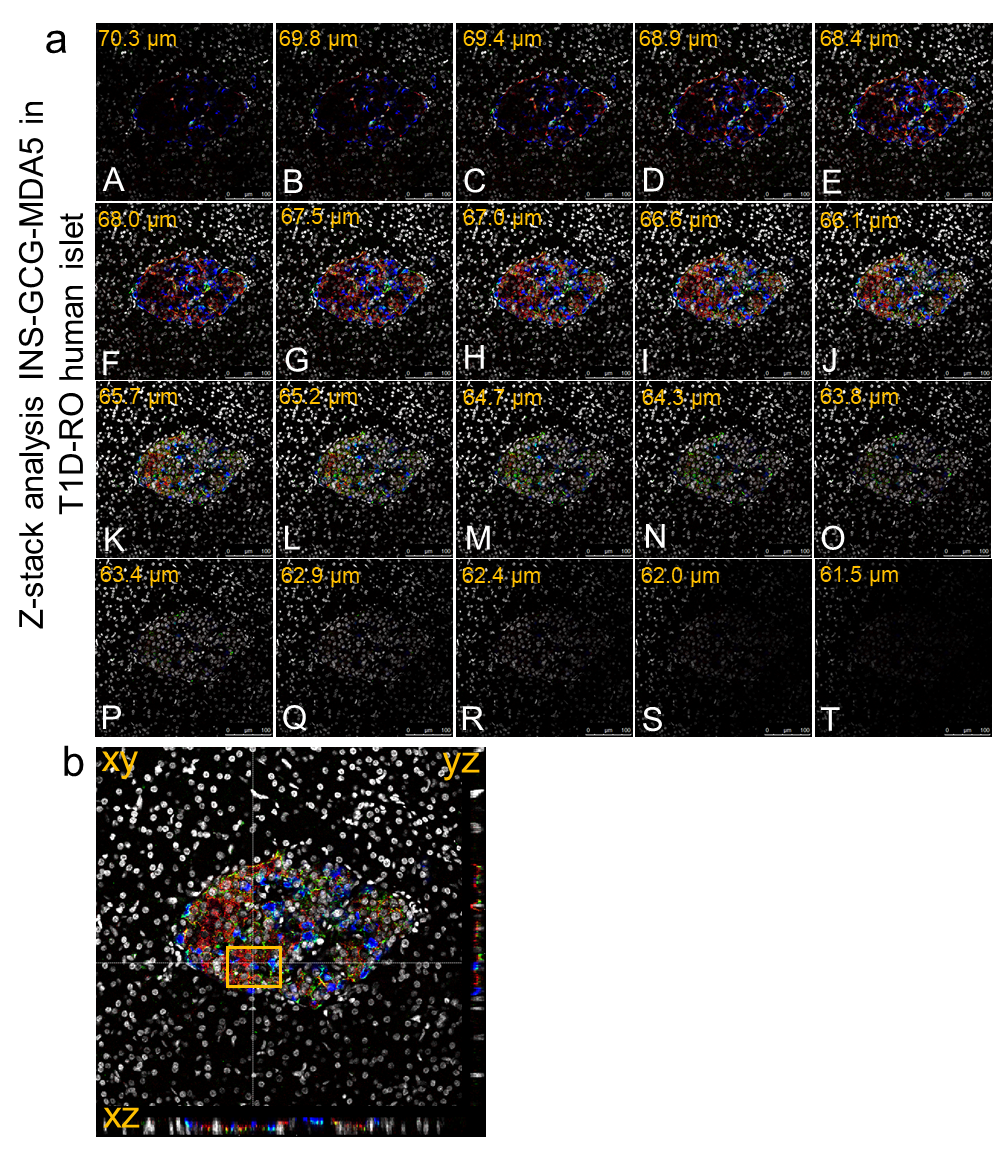

**Supplementary Figure 2. a)** Representative Z-stack analysis of insulin, glucagon and MDA5 using triple immunofluorescence staining of human pancreatic tissue of a T1D recent-onset donor. Images series (panels A to T) show the serial z-stack (n=20 confocal plane) relative to intracellular distribution of insulin (red), glucagon (blue) and MDA5 (green) in a human pancreatic islet of T1D recent-onset donor. **b**) xz and zy imaging deconvolution analysis projection showing intracellular distribution of insulin (red), glucagon (blue) and MDA5 (green) in a human pancreatic islet of T1D recent-onset donor.

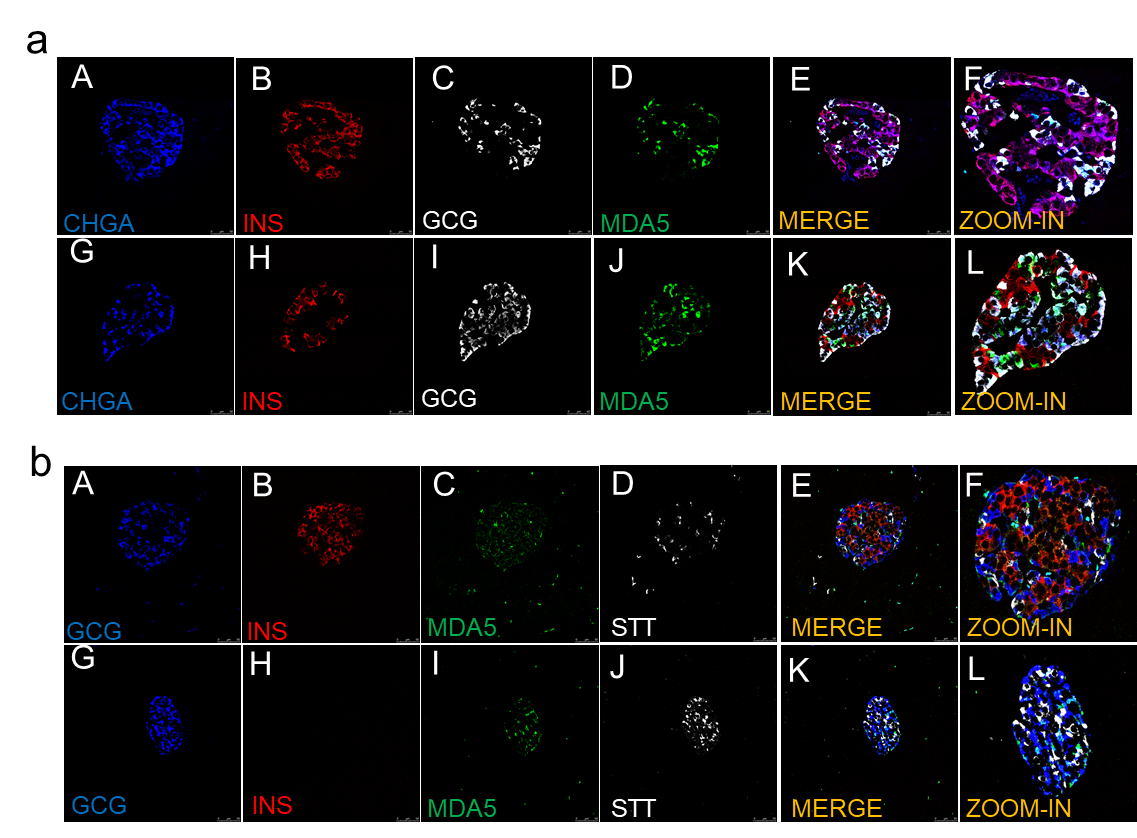

**Supplementary Figure 3. a**) Quadruple immunofluorescence analysis of chromogranin, insulin, glucagon and MDA5 in human pancreatic sections of T1D multiorgan donors. Representative images show fluorescence confocal microscopy imaging analysis of FFPE pancreatic tissue sections stained for chromogranin A (blue, panels A, G), insulin (red, panels B, H), glucagon (white, panel C, I) and MDA5 (green, panels D, J).

**b**) Quadruple immunofluorescence analysis of somatostatin, insulin, glucagon and MDA5 in human pancreatic sections of T1D multiorgan donors. Representative images show fluorescence confocal microscopy imaging analysis of FFPE pancreatic tissue section stained for glucagon (blue, panels A, G), insulin (red, panels B, H), MDA5 (green, panels C, I) and somatostatin (white, panel D, J).

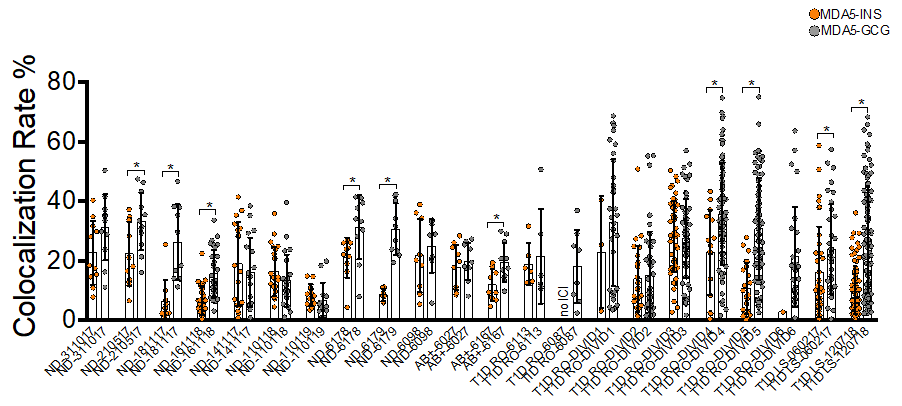

**Supplementary Figure 4.** Dot plot graph showing the colocalization rate (percentage %) between MDA5-insulin (orange circles) and MDA5-glucagon (grey circles) in all cases analysed (ND, Aab^+^, T1D-RO and T1D-LS). Each dot represents an individual islet. Histograms showing mean colocalization value alonside S.D. are reported as well. Statistical analysis was performed using a pairwise comparison (MDA-insulin vs MDA-glucagon for each donor) using a non-parametric Mann-Withney U test (*p value <0.05).

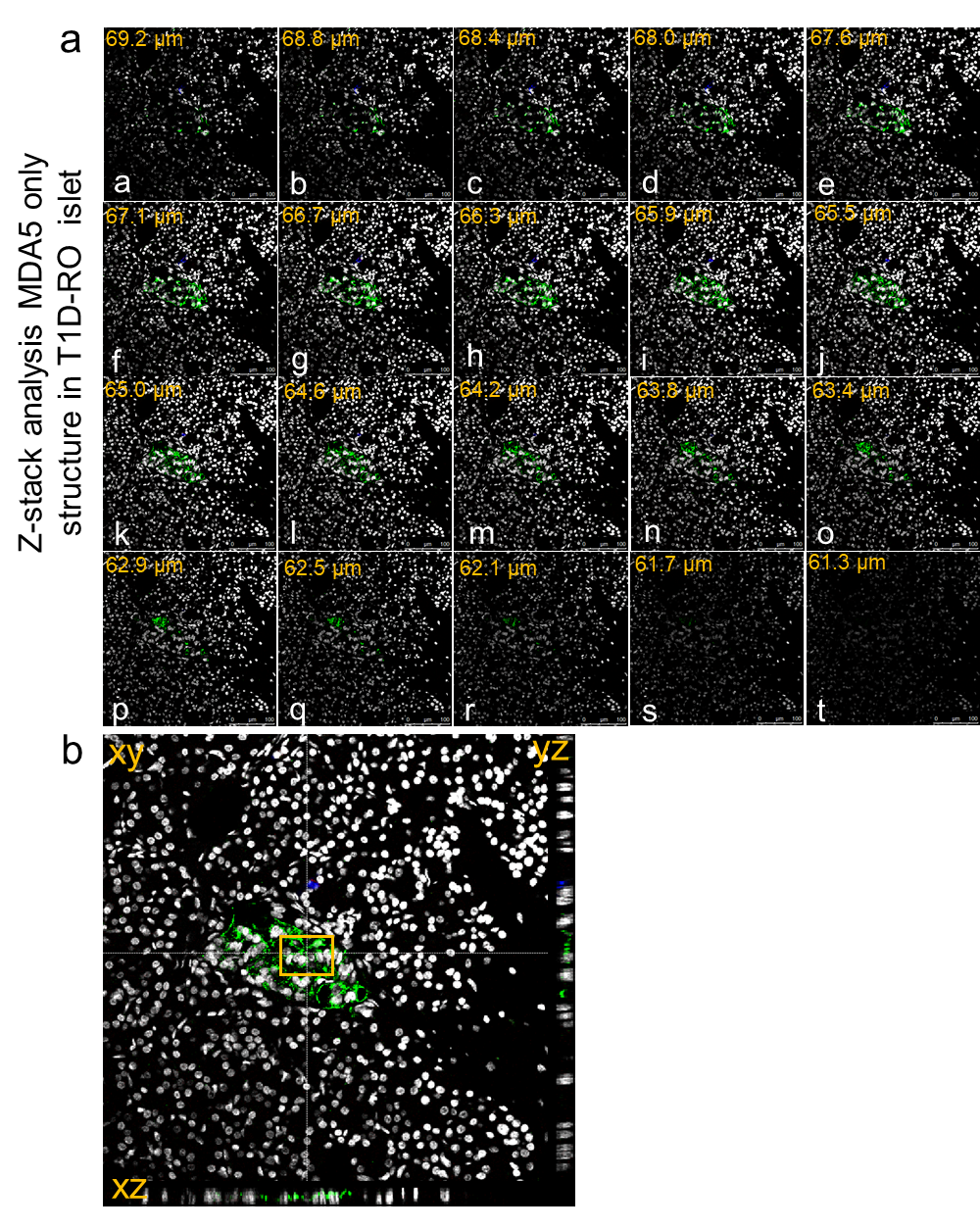

**Supplementary Figure 5**. **a)** Z-stack analysis of MDA5-positive/hormones-negative islets like structure identified using insulin-glucagon-MDA5 triple immunofluorescence staining in human pancreatic tissues of T1D recent-onset donor. Images series (panels A to T) show the serial z-stack (n=20 confocal plane) relative to intracellular distribution of insulin (red), glucagon (blue) and MDA5 (green) in a human pancreatic islet of T1D recent onset donor.

**b)** xz and zy imaging deconvolution analysis projection showing intracellular 3D distribution MDA5 (green) in a MDA5-positive/hormone-negative islets like structure.
